# Transcriptional profiling during infection of potato NLRs and *Phytophthora infestans* effectors using cDNA enrichment sequencing

**DOI:** 10.1101/2024.02.16.580628

**Authors:** Amanpreet Kaur, Vikrant Singh, Stephen Byrne, Miles Armstrong, Thomas Adams, Brian Harrower, Ewen Mullins, Ingo Hein

**Author notes:** Corresponding authors., (I. Hein), (E. Mullins).

## Abstract

An accurate assessment of host and pathogen gene expression during infection is critical for understanding the molecular aspects of host-pathogen interactions. Often, pathogen-derived transcripts are difficult to ascertain at early infection stages owing to the unfavourable transcript representation compared to the host genes. In this study, we compare two sequencing techniques, RNAseq and enrichment sequencing (RenSeq and PenSeq) of cDNA, to investigate gene expression patterns in the doubled monoploid potato (DM) infected with the late blight pathogen *Phytophthora infestans*. Our results reveal distinct advantages of cDNA RenSeq and PenSeq over traditional RNAseq in terms of target gene representation and transcriptional quantification at early infection stages. Throughout the infection time course, cDNA enrichment sequencing enables transcriptomic analyses for more targeted host and pathogen genes. For highly expressed genes that were sampled in parallel by both cDNA enrichment and RNAseq, a high level of concordance in expression profiles is observed, indicative of at least semi-quantitative gene expression representation following enrichment.

## 1. Introduction

The potato late blight pathogen *Phytophthora infestans* causes about 20% of crop losses annually, resulting in a cost of £3.5 billion globally [1]. In the absence of durable resistances, most currently grown commercial potato varieties are vulnerable to the pathogen and often protected through repeated chemical applications, which can exceed 15 sprays per crop annually in conditions that are favourable for pathogen proliferation [2]. Furthermore, the withdrawal of fungicides or the development of fungicide resistance in *P. infestans* can lead to the failure of widely used chemical control measures, which can take a decade to develop and be approved [3]. This has added to the cost and complexity of potato crop protection methods, whilst the cost of chemical-based late blight control has risen by about 25% in developed nations [4]. To identify additional control strategies, a more detailed understanding of the molecular mechanisms involved in *P. infestans* pathogenicity is needed.

*P. infestans* is a hemibiotrophic pathogen that requires living host tissue in the initial biotrophic phase and which progresses to a necrotrophic phase leading to sporulation in a susceptible host [5]. Important features of *P. infestans* infection are haustoria which are indicative of a closely entwined relationship with the host [6]. These specialised invasion organs act as a junction to deliver apoplastic or cytoplasmic effectors which manipulate host metabolism and also suppress host defences [7]. In *P. infestans*, many cytoplasmic effectors have a canonical structure comprising an N-terminal signal peptide, an RxLR motif and a C-terminal effector domain; thus known as RxLRs [8]. As a defence mechanism, plants employ intracellular disease resistance genes that respond to effector function [9]. Known and recognised *P. infestans* RxLRs include, for example, *Avr1, Avr2, Avr3a, Avr3b, Avrblb1, Avrblb2, Avrvnt1, Avramr1*, and *Avramr3* (reviewed in [10]). These, in turn, are recognised by the host genes *R1, R2, R3a, R3b, Rpi-blb1, Rpi-blb2, Rpi-vnt1, Rpi-amr1,* and *Rpi-amr3,* all of which belong to the family of nucleotide-binding, leucine-rich-repeat (NLR) disease resistance genes (reviewed in [10]). In the Solanaceae plants tomato and potato, NLRs are often expressed constitutively at low levels, enabling plants to remain poised for pathogen detection, albeit tissue specific regulatory patterns have been observed [11].

The application of PenSeq to genomic DNA has provided insight into the allelic diversity of *P. infestans* effectors [12] and has also been described for environmental samples [13]. As effectors accounts for < 1% of the *P. infestans* genome, targeting less than 0.5 Mbp of the 240 Mbp *P. infestans* genome, PenSeq reduces the genome complexity and enables evolutionary studies with higher statistical power [12]. However, genomic DNA-based PenSeq does not provide any information about effector expression levels. Therefore cDNA PenSeq was developed, which allowed identification of 47 additional *P. infestans* RxLRs expressed during infection [14].

Similarly, RenSeq, when applied to genomic DNA has proven to be a versatile tool. In potatoes, RenSeq has been used to map and clone new resistances such as *Rpi-ver1*, *Rpi-blb4*, *Rpi-amr1*, *Rpi-amr3i* and *Rpi-amr3* from the wild potato species *S. verrucosum*, *S. bulbocastanum* and *S. americanum,* respectively [15–18]. RenSeq has also been used to track the deployment of functional NLRs in cultivars (dRenSeq; [19]), and to represent full-length NLRs by the development of long-read enrichment sequencing with PacBio, SMRT-RenSeq [20]. Recently, RenSeq-based association studies have been reported in the form of AgRenSeq [21], SMRT-AgRenSeq [22] and SMRT-AgRenSeq-d, to rapidly identify candidate genes associated with elusive resistances in potato cultivars [23]. RenSeq has also been applied to cDNA to provide information about low expressed NLRs which otherwise cannot be identified by RNAseq [24].

Although the role of PenSeq and RenSeq in exploring effector diversity and the identification of novel resistances has been well established, it remains unknown if cDNA enrichment sequencing of effectors and NLRs can provide critical differential expression insight into early infection time points which often remain elusive in standard RNAseq studies. To address this question, we infected microshoots of the *Solanum tuberosum* group Phureja clone DM 1-3 516 R44 with the virulent *P. infestans* isolate W9928C. Extracted cDNA samples were subjected, in parallel, to RNAseq, PenSeq, and RenSeq-base sequencing. This is the first comparative study to establish the quantitative capabilities of RenSeq and PenSeq enrichment techniques.

## 2. Material and Methods

### 2.1. Sample Preparation

Tissue culture plants of the doubled monoploid potato *S*. *tuberosum* group Phureja DM 1-3 516 R44 (commonly known as DM) were maintained on MS20 medium (Murashige and Skoog medium, Duchefa, containing 20 g/L sucrose; [25]. Plants were kept in a growth room at a light intensity of 110 µmol m^-2^s^-1^, a temperature of 18 ± 2 °C, and a photoperiod of 16/8h light/dark [25].

For the experiment, healthy three-week-old DM plantlets with fully expanded leaves were selected. *In vitro* shoots along with roots were gently removed from the media and dipped for one minute in a zoospore suspension of *P. infestans* isolate W9928C adjusted to 4 x 10^6^ spores/mL [26]. Dip-inoculated microshoots were gently blotted on sterile paper towels and again planted in fresh MS20 media in vented tissue culture grade glass containers (Generon, U.K.). The infected plants were kept in darkness for 16 hours and then incubated under the growth conditions mentioned above. The disease severity was recorded by counting the number of leaves showing disease symptom in 24-hour intervals. The leaf samples from four independent replicates were collected after 0, 24, 48, 72 hours post infection (hpi) and immediately immersed in liquid Nitrogen before storing at -70 °C for further processing (Supplementary Figure 1). Additional samples were incubated for 96 hours for visual inspection only.

### 2.2. RNA isolation and cDNA synthesis

From each individual replicate taken at 0, 24, 48 and 72 hpi, leaf samples were crushed to a fine powder and RNA was extracted combining TRI reagent (Sigma-Aldrich) and RNeasy kit (Qiagen, Germany) protocols. Specifically, 400 mg of ground sample was resuspended in 2 mL of TRI reagent and vortexed after addition of 10 µL β-mercaptoethanol. The slurry was left to stand at room temperature for 5 minutes before centrifugation at 10,000g for 10 min at 4 °C. Chloroform was added to the supernatant (0.2 mL per mL), kept at room temperature for 5 minutes, and centrifuged. Isopropanol (0.5 mL per mL) was added to the aqueous phase and the solution was transferred to a QIA RNAeasy spin column for binding and washing in RPE buffer twice. RNA was eluted in RNAse free water (50 µL), and the integrity was checked using a Bioanalyzer 2100 (Agilent).

### 2.3. Sequencing

A pipeline was developed for parallel processing the samples for RNAseq and enrichment sequencing (PenSeq and RenSeq; Supplementary Figure 1). Quality checked RNA (RIN value ≥8) was divided into two parts; one part was processed at the James Hutton Institute’s Genomics facility for generating RNA sequencing libraries using the stranded Illumina mRNA Prep kit, and Integrated DNA technology (IDT) RNA unique dual UD Indices (Illumina) as recommended, with 100 ng total RNA per sample. Libraries were quality checked on a fluorimeter (Qubit) and on a Bioanalyzer 2100 prior to pooling equimolar amounts before sequencing. Sequencing was conducted on a NextSeq 2000 (Illumina) sequencer at loading concentration of 750 pM using a P3 200 kit, generating paired-end 100 bp reads.

In parallel, second strand cDNA was synthesised from the RNA using both oligo(dT) and random hexamer primers following the manufacturer’s guideline (Superscript IV, Thermo). Individually indexed libraries were prepared for PenSeq and RenSeq by combining equimolar amounts of cDNA from each time point and replicate. Enrichment sequencing was conducted using the myBaits v5.02 protocol in combination with the PenSeq bait library D10251HRPen and RenSeq library vs5 D10580RnSq5, respectively at Arbor BioSciences, MI, USA and the captures were pooled for sequencing on an Illumina NovaSeq 6000 platform on a partial S4 PE150 lane.

### 2.4. Data analysis

Reads obtained from RNAseq and cDNA enrichment sequencing (RenSeq/PenSeq) were subjected to quality control and adapter trimming using fastp [27] at 99% base call accuracy (default settings allowing phred quality 20). Trimmed reads were mapped to the host (*S. tuberosum* group Phureja clone DM1-3 516 R44 v6.1) and pathogen (*P. infestans* T30-4) reference genomes using a bowtie based splice aware mapper, HISAT2, under stringent conditions allowing 1% mismatch rate (--score-min L,-0.06,-0.06; [28]). The reads were also quantified against reference transcriptomes with Salmon running in alignment mode for quantification of the bam files. The quantification files were analysed for differential gene expression using the 3D RNAseq App [29]. Read counts and transcript per million reads (TPMs) were established using tximport R package version 1.10.0 and lengthScaledTPM method [30] with inputs of transcript quantifications from Salmon [31]. Low expressed transcripts and genes were filtered based on analysing the data mean-variance trend. The Trimmed Means of M values (TMM) method was used to normalise the gene and transcript read counts to log2 CPM [32] and a principal component analysis (PCA) was performed to identify any batch effects. The limma R package was used for gene and transcript expression comparison [33]; [34]. The log2 fold change (FC) of gene and transcript abundance was calculated and a *t*-test was used to determine significance of expression changes. *P*-values of multiple testing were adjusted with the Benjamini-Hochberg (BH) procedure to correct false discovery rate (FDR ≤ 0.05; [35]), according to the built-in protocol of 3D-RNAseq app [29]. A gene was considered significantly differential expressed if it had an adjusted *p*-value of < 0.01 and log2 FC ≥ 1.

To filter NLRs from the expressed genes, NLR annotation was conducted by running NLR tracker [36] on coding sequence (CDS) of DM v6.1. The sequences of retrieved NLRs were subjected to a BLAST analysis with the previously reported 755 NLRs by [37]. Only 100% matching sequences with available reference gene IDs were retained for the expression analysis yielding 583 NLRs.

To calculate the sequencing coverage of NLRs and pathogen targets (RxLR and non-RxLR), bed files generated from genome assemblies of DM v4.03 [37] and *P. infestans* T30-4 [12] with coordinates of 755 NLRs and 579 effectors (RxLRs and non-RxLRs) were used as a reference. Sequencing depth and coverage of target genes were calculated using bedtools v2.31.0 and compared between RNAseq and cDNA enrichment (RenSeq and PenSeq). Data were further processed to establish correlations among the datasets and time points using Pearson’s correlation. Before analysing the TPM values were transformed using log10 (TPM+1).

## 3. Results

Microshoots of DM infected with *P. infestans* isolate W9928C showed progressive late blight disease symptoms (Figure 1). Visible disease symptoms developed at 48 hours post infection (hpi) which coincides with the transition between the biotrophic and necrotrophic phases. At 72 hpi, most of the leaves showed lesions which proceeded to complete necrosis at 96 hpi. Thus, 96 hpi samples were excluded from the sequence-based analyses. Leaf samples from the infected plants were collected at 0, 24, 48, and 72 hpi for RNAseq and RenSeq/PenSeq-based cDNA enrichment sequencing, respectively. Following Illumina sequencing, 1.13 billion paired end reads (100 bp) showing an overall Q30 > 94% were obtained for RNAseq (Table S1). For PenSeq and RenSeq-based enrichment sequencing, a total of 369 million reads passed the filter (PF; Q30) with read counts ranging between 19 million to 28 million per sample (Table S1).

**Figure 1:**
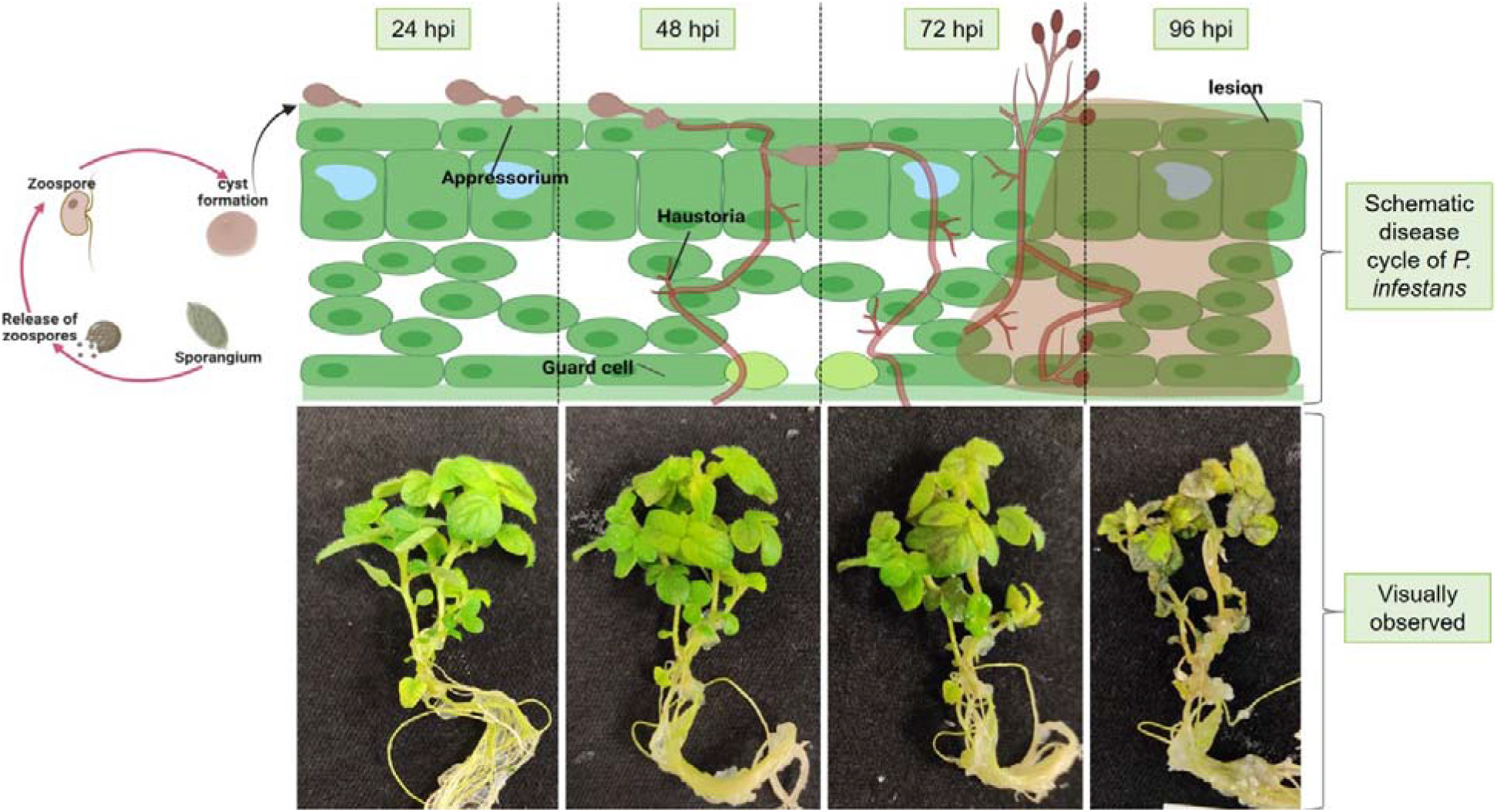
Diagrammatic representation of various stages of *Phytophthora infestans* life cycles and disease progression on *in vitro Solanum tuberosum* group Phureja DM 1-3 516 R44 (commonly known as DM) plants.

Following QC and adaptor trimming, more than 832 million reads from RNAseq and over 363 million reads from enrichment sequencing were retained (Table S1). These reads were mapped to the potato (*S*. *tuberosum* group Phureja DM 1-3 516 R44 v6.1) and *P. infestans* (T30-4) reference genomes at 1% mismatch rates. For RNAseq, 749 million reads out of the 832 million reads (89.98%) mapped to the potato reference genome whereas only 1.6 million (0.2%) mapped to *P. infestans* reference genome (Table 1). For the indexed RenSeq and PenSeq enrichment sequencing, a total of 363 million reads were mapped to reference genomes, out of which 280 million (76.99%) mapped to potato (RenSeq) and 8.3 million (2.28%) mapped to *P. infestans* (PenSeq). Importantly, in the RNAseq analysis the percentage of pathogen derived reads did not increase during the infection time course whereas the PenSeq analysis displayed a steady increase of detectable pathogen transcripts from 1.04% at 0 hpi, 1.47% at 24 hpi, 2.52% at 48 hpi and 4.04% at 72 hpi (Table 1).

**Table 1:** Number of RNAseq, RenSeq and PenSeq reads mapped to the host and pathogen genomes. *Solanum tuberosum* group Phureja DM 1-3 516 R44-v6.1 (DM v6.1) and *Phytophthora infestans* strain T30-4 genomes were used as references and reads obtained at different infection timepoints were mapped using HISAT2 (Kim et al., 2019).

| Sequencing techniques | Total reads | Reads mapped to DM v6.1 | % reads mapped to DM v6.1 | Reads mapped to *P. infestans* T30-4 | % reads mapped to *P. infestans* T30-4 |
| --- | --- | --- | --- | --- | --- |
| **RNAseq** |  |  |  |  |  |
| **Hours post infection** |  |  |  |  |  |
| 0 | 192,017,649 | 190,972,662 | 99.4250 | 527,619 | 0.27 |
| 24 | 225,017,104 | 222,731,572 | 98.9925 | 258,190 | 0.11 |
| 48 | 214,620,790 | 205,952,655 | 95.9950 | 429,980 | 0.20 |
| 72 | 201,262,977 | 129,825,714 | 64.9825 | 483,111 | 0.24 |
| **Total** | **832,918,520** | **749,482,603** | **89.9800\*** | **1,698,900** | **0.21\*** |
| **RenSeq** |  |  |  |  |  |
| 0 | 95,324,230 | 80,302,072 | 84.2400 |  |  |
| 24 | 85,202,507 | 73,723,461 | 86.5300 |  |  |
| 48 | 88,339,763 | 75,416,568 | 85.3700 |  |  |
| 72 | 95,095,897 | 50,796,658 | 53.4200 |  |  |
| **Total** | **363,962,397** | **280,238,759** | **76.9900\*** |  |  |
| **PenSeq** |  |  |  |  |  |
| 0 | 95,324,230 |  |  | 992,390 | 1.04 |
| 24 | 85,202,507 |  |  | 1,248,629 | 1.47 |
| 48 | 88,339,763 |  |  | 2,227,074 | 2.52 |
| 72 | 95,095,897 |  |  | 3,842,593 | 4.04 |
| **Total** | **363,962,397** |  |  | **8,310,686** | **2.27\*** |

\*average across the time points

### 3.1. Targeted genes are highly represented by RenSeq and PenSeq reads

The mapped RNAseq, RenSeq and PenSeq reads were filtered for the 579 target genes in *P. infestans* (including 438 RxLR and 141 non-RxLR effectors; [12]) and 755 potato NLRs [37] using bedtools. The number of *P. infestans* targets and NLRs expressed in RNAseq and enrichment (RenSeq and PenSeq) datasets were evaluated (Table 2). Out of a total of 579 *P. infestans* effectors, an average of 316 genes (289 at 0 hpi, 332 at 24 hpi, 331 at 48 hpi, and 312 at 72 hpi) were represented by cDNA-PenSeq reads indicative of their expression, whereas an average of 167.25 *P. infestans* target genes (146 at 0 hpi, 181 at 24 hpi, 192 at 48 hpi, and 150 at 72 hpi) were detected by RNAseq reads (Table 2, Figure 2A). Thus, a two-fold increase in the number of expressed target genes was observed in PenSeq in comparison with RNAseq. Importantly, all genes identified by RNAseq were also represented by cDNA-based PenSeq.

**Figure 2:**
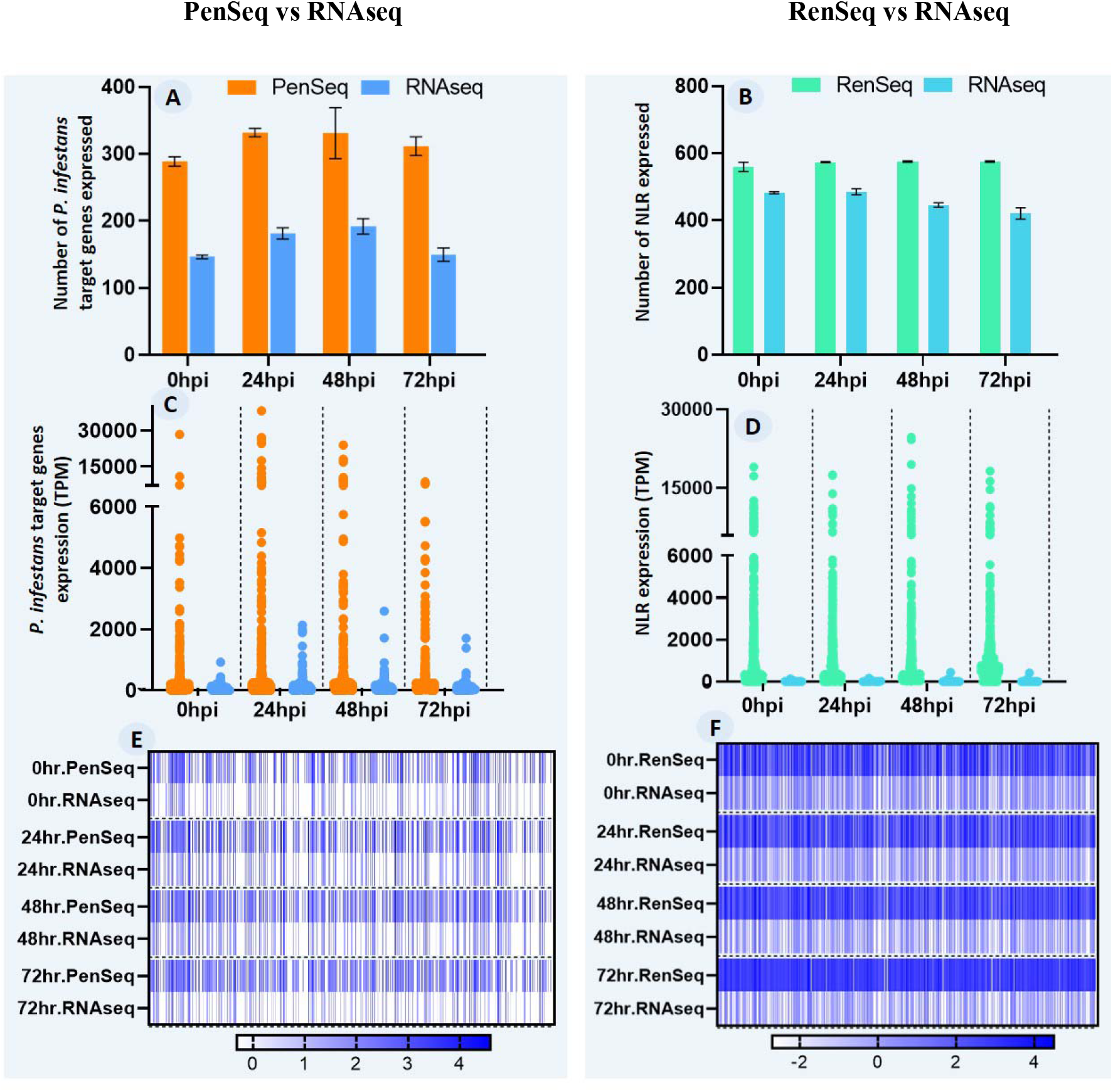
A comparison between RNAseq and PenSeq/RenSeq enrichment sequencing for the expression of pathogen effectors (RxLR and non-RxLR effectors) and NLRs at different time points of infection. **A)** Number of effectors expressed in RNAseq and PenSeq datasets following mapping to the *Phytophthora infestans* transcriptome at high stringency allowing a 1% mismatch rate **B)** Number of NLRs expressed in RNAseq and RenSeq datasets following mapped to *Solanum tuberosum* group Phureja DM 1-3 516 R44-v6.1 (DM v6.1) transcriptome at a 1% mismatch rate **C-D)** A comparison between the expression values (Lengthscaled Transcripts Per Million; TPM values) of effectors and NLRs between RNAseq and enrichment techniques **E-F)** Heatmap comparing effector and NLR expressions between RNAseq, PenSeq and RenSeq. TPM values were normalized to log10(TPM+1) before generating heatmaps.

**Table 2:** A comparison of expression values (Transcripts Per Million; TPM) of effectors (RxLRs and non-RxLRs) determined using PenSeq and RNAseq data. The mean of TPM values across the four replicates was utilised for the comparison.

| Hours post infection (hpi) | PenSeq |  | RNAseq |  |
| --- | --- | --- | --- | --- |
|  | Total effectors expressed (Mean) | Mean number of expressed effectors with TPM > 100 | Total effectors expressed (Mean) | Mean number of expressed effectors with TPM > 100 |
| 0 | 288.75 | 174.25 | 146.25 | 25.75 |
| 24 | 331.75 | 226.25 | 181.25 | 57.00 |
| 48 | 331.00 | 214.75 | 192.00 | 40.50 |
| 72 | 311.50 | 205.00 | 149.50 | 21.25 |
| Average Number of expressed effectors | 315.75 | 205.06 | 167.25 | 36.13 |
| Percent effectors with TPM>100 | 64.94 |  | 21.59 |  |

To evaluate NLR expression, the reads were mapped to the latest transcript assembly of potato (DM v6.1). NLR sequences identified using NLR tracker were subjected to BLAST using the previously reported 755 NLRs in DM v4.3 [37] and we could determine the position of 583 NLRs with existing reference transcript IDs. Of these 583 indexed NLRs, an average of 571 NLRs was detectable in cDNA RenSeq across all the infection points whereas only 429 NLRs were expressed according to RNAseq data (Table 3, Figure 2B). Critically, all genes identified as expressed by RNAseq were also detectable by cDNA RenSeq.

**Table 3:**
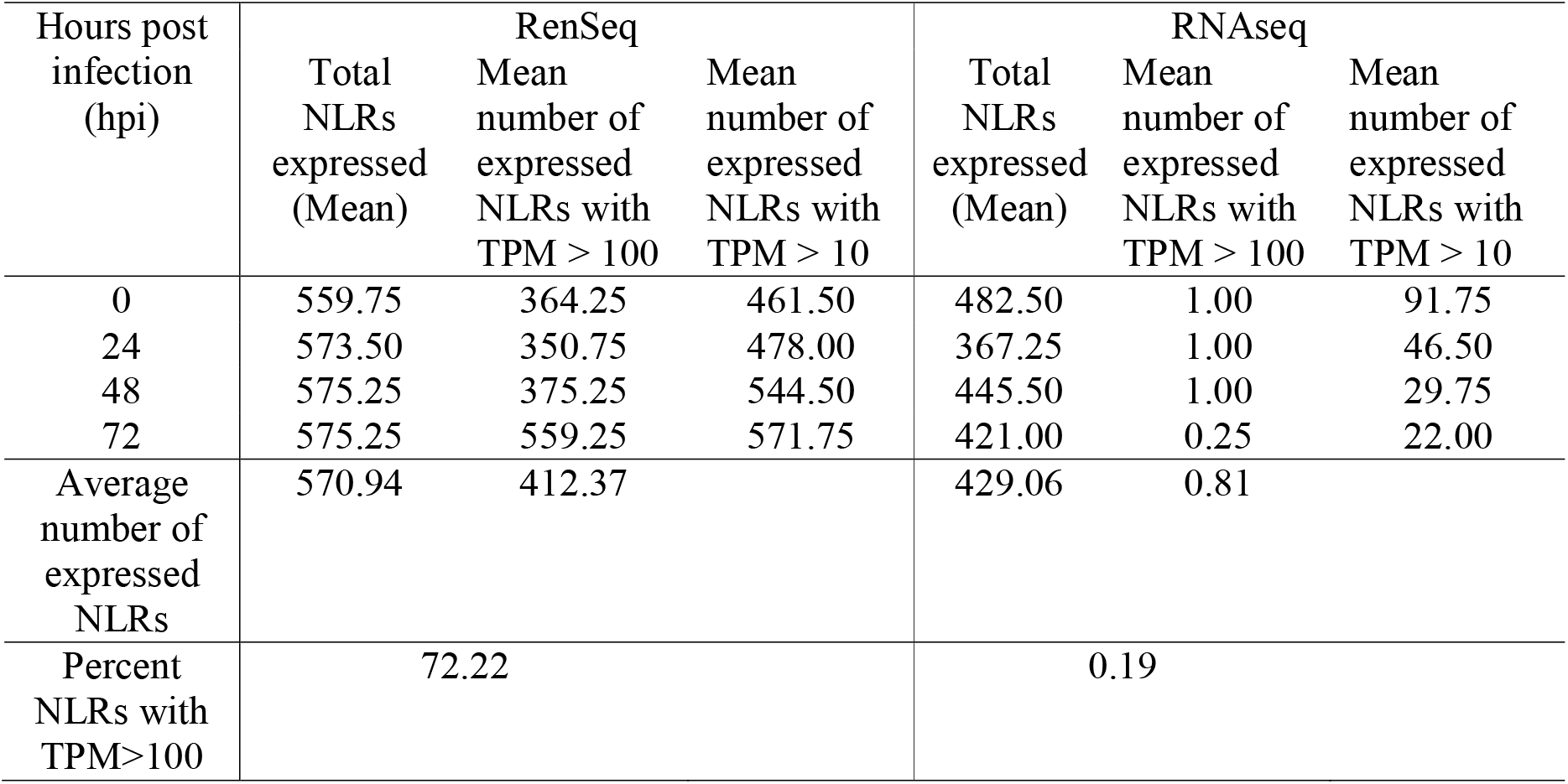
A comparison of expression values (Transcripts Per Million; TPM) of NLRs determined using RenSeq and RNAseq data. A mean TPM value of the four replicates was calculated for the comparison. Expression was calculated only for the 583 NLRs for which reference gene IDs are available in the latest potato transcriptome version (v6.1).

### 3.2. Enriched mapping resulted in higher expression values of the target genes

The length scaled transcript per million (TPM) values of the target genes in *P. infestans* and potato were highest in the enriched datasets (PenSeq and RenSeq) in comparison to RNAseq (Figure 2C-F). For *P. infestans*, an average of 64.94% of expressed effectors (RxLRs and non-RxLRs) yielded TPM values greater than 100 in the PenSeq dataset, whereas this was limited to 21.59% in RNAseq derived reads (Figure 2C, Table 2, Table S2). A similar trend was observed for potato, where an average of 72.22% of expressed NLRs have TPM values greater than 100 in the RenSeq-based analysis. In contrast, only 0.19% expressed NLRs in RNAseq reads showed TPM values greater than 100 (Figure 2D, Table 3, Table S3).

3.3. High correlation between RNAseq and enrichment datasets validates concordance between RNAseq and enrichment techniques

To assess if cDNA targeted enrichment sequencing retains quantitative transcriptional properties, expression patterns of genes that were detectable in RNAseq and PenSeq/RenSeq-derived datasets were compared. Across the infection time points, 579 pathogen effector genes and 583 host NLRs were used to calculate a Pearson correlation coefficient (Supplementary Figure 2). The computed correlation coefficient for pathogen target genes expressed in four replicates was in the range of 0.84[–0.98 and 0.78[–0.87 within and between the PenSeq and RNAseq datasets, respectively (Supplementary Figure 2A, Table S4). A similar high correlation (0.77–0.98) of NLRs expressed in the replicates and within as well as between both techniques (targeted enrichment and RNAseq) was observed (Supplementary Figure 2B, Table S4). This shows a strong consensus between the replicates as well as the techniques.

A quantitative comparison of pathogen target gene expression levels [log10 (TPM+1)] between PenSeq and RNAseq indicated a significant correlation among the techniques (Supplementary Figure 2C). It was interesting to find that 149 of the target genes were expressed only in the PenSeq data and not detectable by RNAseq. This demonstrates a distinct advantage of enrichment over RNAseq. A comparison between NLR expression levels between RenSeq and RNAseq showed a similar trend, where 142 additional NLRs were detected only by RenSeq (Supplementary Figure 2D).

The expression profiles of pathogen target genes and NLRs as detected by RenSeq and PenSeq were subjected to hierarchical clustering to group the genes into clusters with similar expression patterns (Figure 3). The expression values of the selected genes from RenSeq and PenSeq when compared with RNAseq showed similar expression patterns (Figure 3). To highlight our observations, the known effectors linked with the biotrophic phase [PITG_04314 (*PexRD24*), PITG_14371 (*Avr3a*), PITG_20300 (*Avrblb2*), PITG_21388 (*Avrblb1*), PITG_03192, PITG_04089] and the closest homologs of functional resistance genes (Soltu.DM.04G006260 [*Rpi-blb3*], Soltu.DM.04G006450 [*R2*], Soltu.DM.04G008180 [*Rpi-amr3*], Soltu.DM.05G005840 [*R1*], Soltu.DM.06G001240 [*Rpi-blb2*], Soltu.DM.08G020910 [*Rpi-blb1*], Soltu.DM.09G029510 [*Rpi-vnt1*], Soltu.DM.09G029560 [*R9a*], Soltu.DM.09G030280, [*R8*] Soltu.DM.11G024990 [*R3a*]) were compared in their expression values in PenSeq, RenSeq and RNAseq datasets. The expression trends of the genes at different infection time points were very similar between the techniques, although the genes from cDNA enrichment showed higher values (Figure 3).

**Figure 3:**
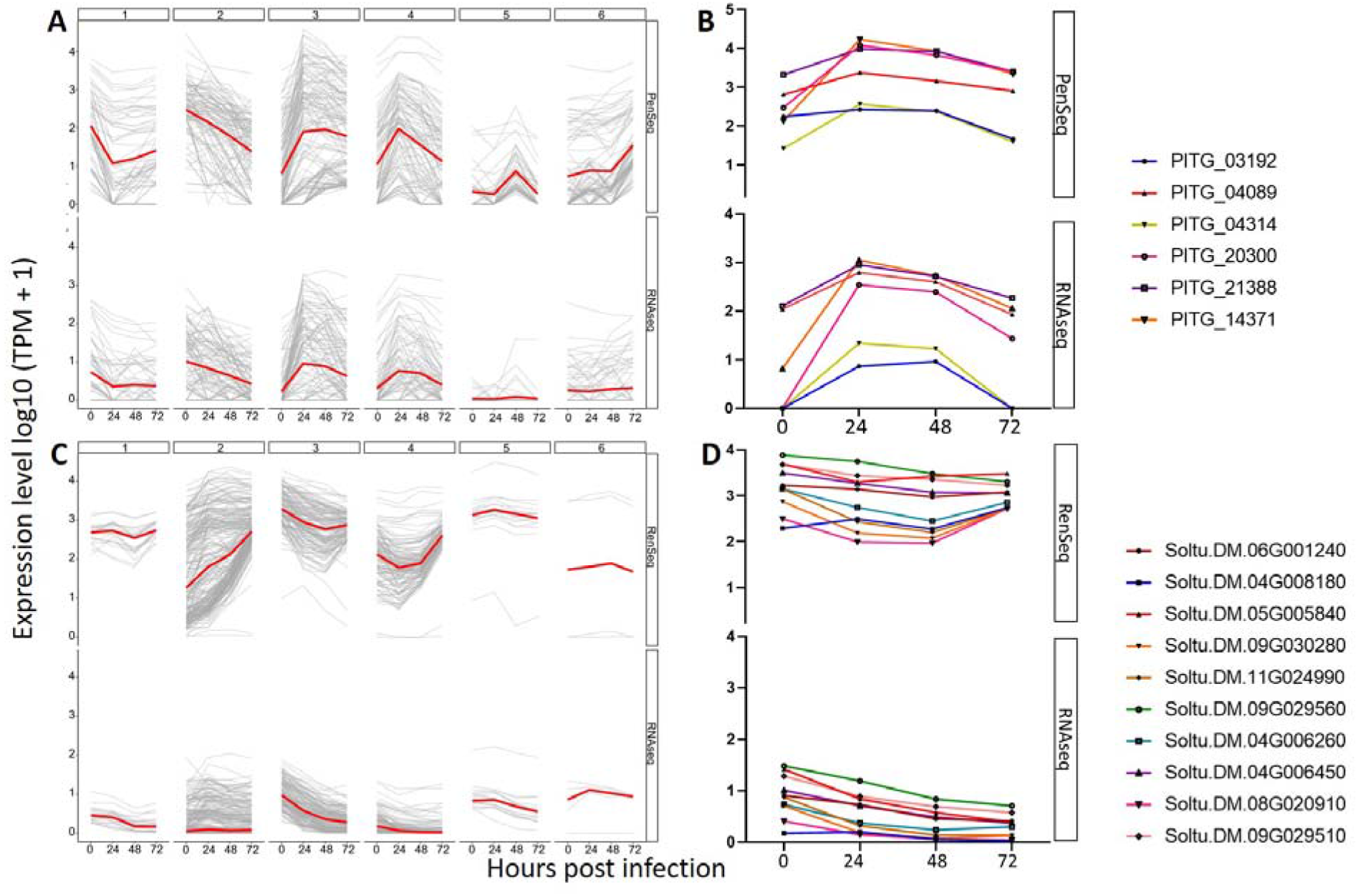
A comparison between expression profiles of effectors (RxLR and non-RxLR) and NLR genes **A)** Expression patterns of effectors expressed in PenSeq and RNAseq datasets **B)** A subset of effectors linked with biotrophic phase of infection was plotted to compare the expression levels **C)** Expression patterns of NLRs expressed in RenSeq and RNAseq datasets **D)** A subset of NLRs (homologs of commercially important *R* genes) was plotted. Transcripts Per Million (TPM) values were transformed to log10(TPM+1) before plotting.

To further validate our observations, previously transcriptionally characterised *P. infestans* marker genes were specifically queried in the data sets. These include the cellulose synthase gene (*CesA4*; PITG_16984) associated with cyst germination and appressorium formation [38], haustorial membrane protein (*HMP1*; PITG_00375) associated with biotrophy [39], and the Necrosis inducing Phytophthora Protein 1 (*NPP1*; PITG_00957), a necrotrophy marker which shows a transition between the biotrophic and necrotrophic phases of infection ([40]; Supplementary figure 3). For the *CesA4* gene (PITG_16984), expression was highest at 24 hpi with a TPM value of 865.17 in the case of PenSeq and 46.93 in RNAseq (Table S2). The TPM values decrease to 620.17 at 48 hpi in the case of PenSeq and to 28.68 in RNAseq. These observations are in accordance with [38] which showed that there is an increase in *CesA* gene expression during the initial phases of infection. For *HMP1*, cDNA PenSeq expression was the highest at 24 hpi with a TPM value of 1,335.51. This decreased to 754.01 at 48 hpi and to 531.56 at 72 hpi (Table S2). The expression profile of *HMP1*, as detected by PenSeq, coincides with the results of the RNAseq analysis where a decrease in the TPM value from 85.22 at 24 hpi to 34.95 at 72 hpi was observed (Table S2). For *NPP1*, PenSeq and RNAseq showed that the expression declined after the onset of the necrotrophic phase, which in our experiment occurred at around 48 hpi. These observations are in line with previous studies which also showed that the expression level of *NPP1* is highest at the beginning of necrotrophic phase [41]. As the difference between the expression values was very high, the TPM values were transformed using log10 (TPM+1) for plotting the graphs to compare gene expression between PenSeq and RNAseq (Supplementary Figure 3).

Overall, these results show that enrichment techniques retain the quantitative expression nature in the data and allow the detection of a greater number of low abundant transcripts compared to RNAseq. Not surprisingly, where genes were only detectable at low expression levels in the RNAseq dataset, the correlation to cDNA enriched profiles was less robust.

As mentioned previously, host and pathogen genes that were represented in the RenSeq and PenSeq bait libraries and identified as expressed by normal RNAseq, were also identified by RenSeq and PenSeq. However, in terms of differential expression, PenSeq identified almost twice as many differentially expressed effectors in comparison to RNAseq in all time points (Figure 4A, Table S5). A modulation of expression of 187, 201, and 159 pathogen target genes (average of 4 replicates) at 24 hpi, 48hpi, and 72 hpi, was identified by PenSeq, while RNAseq identified 87, 84, and 66 target genes only. Critically, most of the differentially expressed genes (DEGs) identified by RNAseq were also identified by PenSeq and for 99.64% of the genes (upregulated or downregulated) the directions of expression changes were identical (Table S5). Since PenSeq was able to identify 43.33% more DEGs in comparison to RNAseq, an average overlap of 39.73% was found between the two techniques (Table 4, Figure 4B).

**Figure 4:**
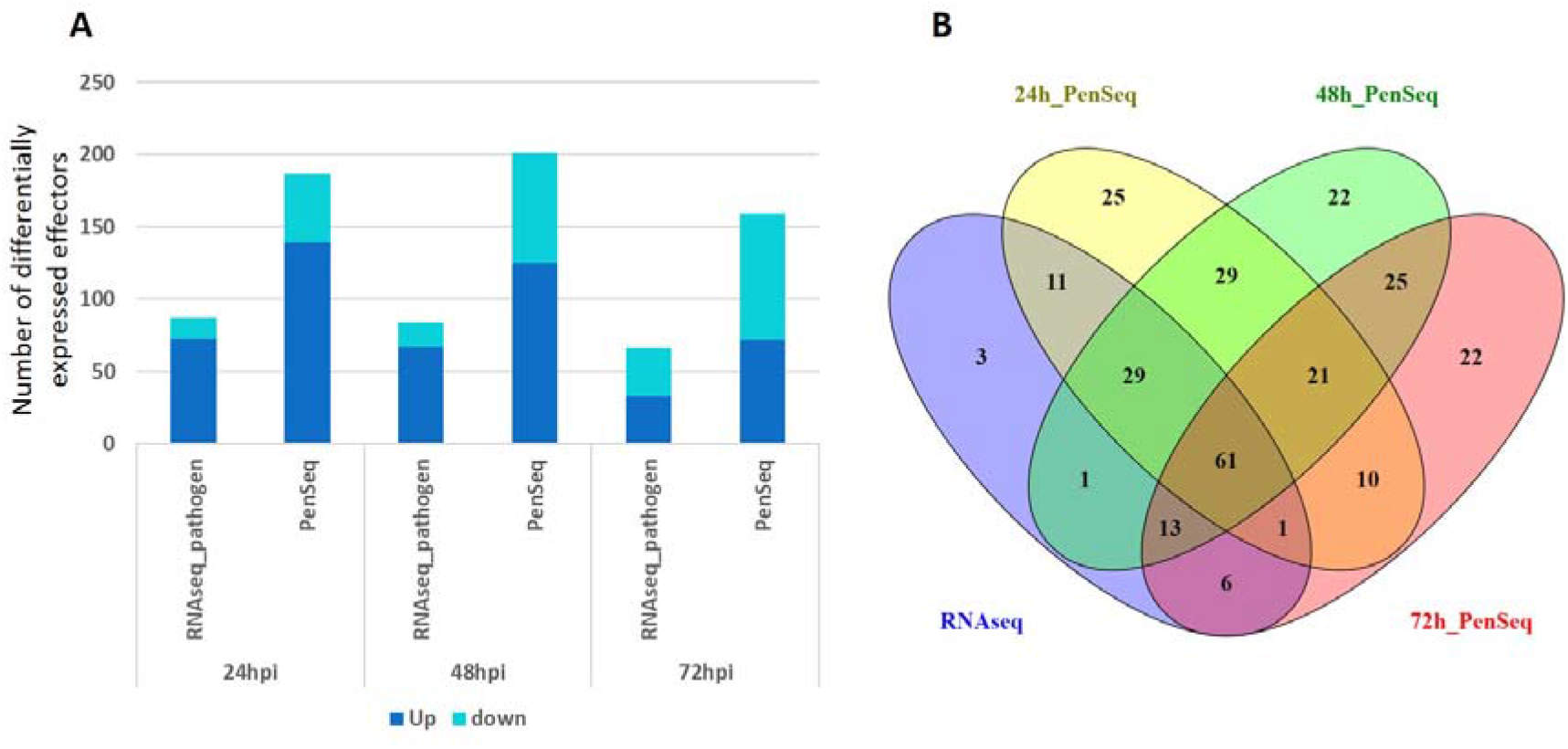
A comparison between differential expression of effectors (RxLR and non-RxLR) between RNAseq and PenSeq. The log2 fold change (FC) of effector abundance was calculated. A *t*-test was used to determine significance of the expression changes. A gene was significantly differentially expressed if it had an adjusted *p*-value < 0.01 and log2 FC ≥ 1. **A)** Variation in the number of effectors expressed significantly between the RNAseq and PenSeq datasets **B)** An overlap between differential effector expression identified by target enrichment (PenSeq) during different infection time points (24, 48, and 72 hours post infection; hpi), and differentially expressed effectors identified by RNAseq throughout infection was observed as a venn diagram.

**Table 4:** Concordance of differentially expressed (DE) pathogen target genes (RxLR and non-RxLRs) between RNAseq and PenSeq approaches.

| Hours post infection (hpi) | DE Pathogen target genes identified in PenSeq | Overlapping DEGs with RNASeq | % of DEGs overlap |
| --- | --- | --- | --- |
| 24 | 187.00 | 83.00 | 44.39 |
| 48 | 201.00 | 77.00 | 38.31 |
| 72 | 159.00 | 58.00 | 36.48 |
| Average | 182.33 | 72.67 | 39.73 |

Similarly, RenSeq identified 163, 301 and 355 differentially expressed NLRs at 24 hpi, 48 hpi and 72 hpi, whereas RNAseq identified 192, 228 and 251 differentially expressed NLRs at these time points (Figure 5A, Table S6). In total, 277 NLRs with significantly modulated expression were identified by RNAseq and the RenSeq datasets with expression concordance. Further, 207 and 43 NLRs were predicted to be differentially expressed exclusively by RenSeq and RNAseq respectively (Table 5, Figure 5B).

**Figure 5:**
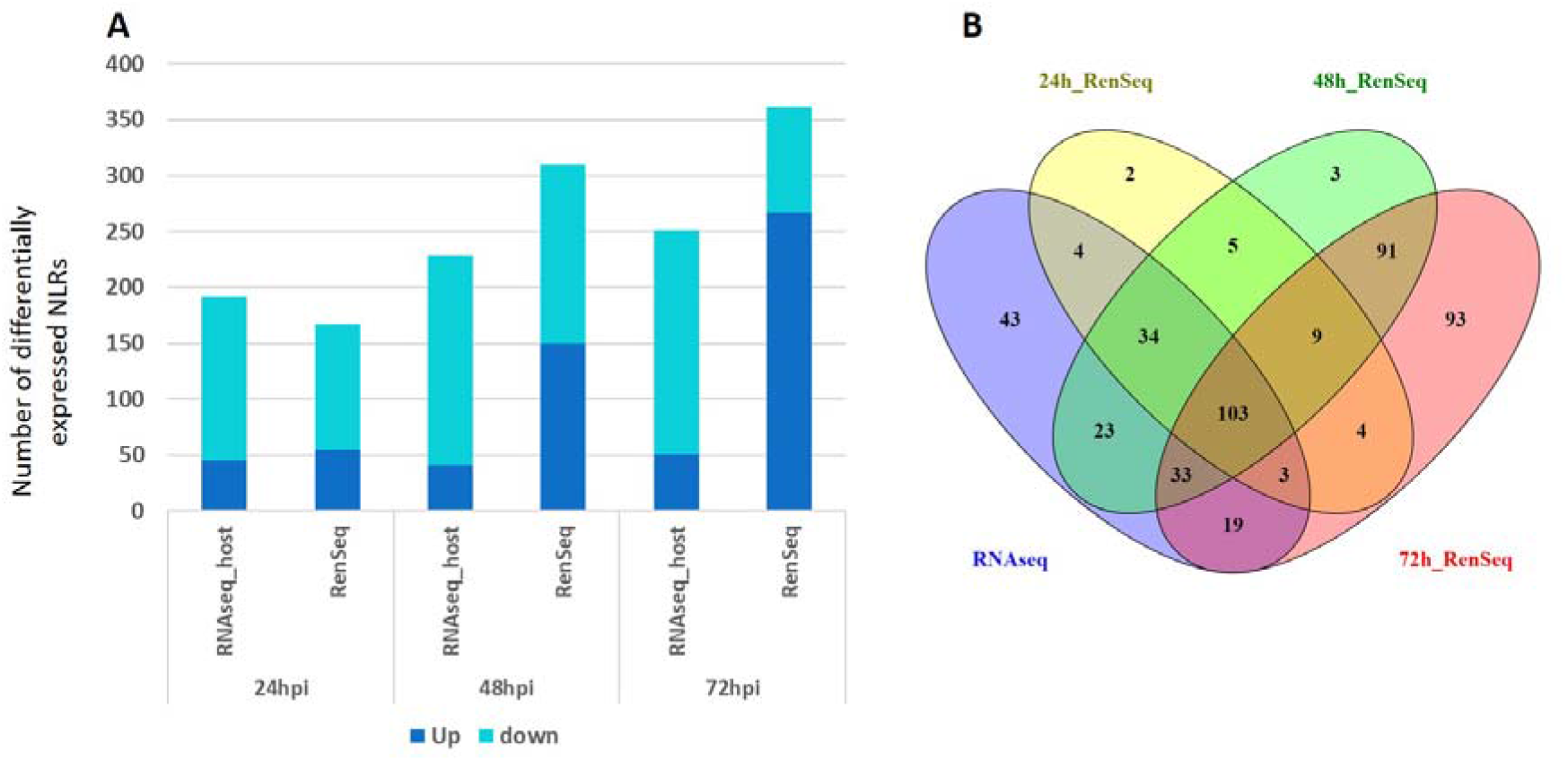
A comparison between differential expression of NLRs between RNAseq and RenSeq. The log2 fold change (FC) of NLR abundance were calculated. A *t*-test was used to determine significance of the expression changes. A gene was significantly differentially expressed if it had an adjusted *p*-value < 0.01 and log2 FC ≥ 1. **A)** Variation in the number of NLRs expressed significantly between the RNAseq and RenSeq datasets **B)** An overlap between differential NLR expression identified by target enrichment (RenSeq) during different infection time points (24, 48, and 72 hours post infection; hpi), and differentially expressed NLRs identified by RNAseq throughout infection was observed as a venn diagram.

**Table 5:** Concordance of differentially expressed (DE) NLRs between RNAseq and RenSeq approaches.

| Hours post infection (hpi) | DE NLRs identified in RenSeq | Overlapping DEGs with RNASeq | % of DEGs overlap |
| --- | --- | --- | --- |
| 24 | 163.00 | 138.00 | 84.66 |
| 48 | 301.00 | 183.00 | 60.80 |
| 72 | 355.00 | 146.00 | 41.13 |
| Average | 273.00 | 155.67 | 62.19 |

### 3.4. cDNA enrichments yield higher target gene coverage

Target gene sequence representation was calculated for the expressed effectors and NLRs as a percentage of gene length covered by the reads (Supplementary Figure 4). In line with the dRenSeq-type analyses, cDNA-derived RenSeq and PenSeq reads showed the highest average number of target genes with 100% coverage in comparison to RNAseq reads (Table 6-7). During infection, out of 288.75, 331.75, 331 and 311.5 (average of 4 replicates) effectors expressed at 0 hpi, 24 hpi, 48 hpi and 72 hpi; a complete (100%) coverage was observed for 99, 132.25, 100.25 and 58 effectors (average of 4 replicates) in the PenSeq data, respectively. This number was limited to 8.75, 15.25, 19.25 and 12.25 effectors (average of 4 replicates) with 100% coverage at various infection time points for RNAseq (Table 6). Similarly, RenSeq data showed 100% coverage in almost twice the number of NLRs detectable by RNAseq during all the time points (Table 7). Complete coverage was observed for 594.75, 606.75, 666.75 and 690.25 NLRs expressed (average of 4 replicates) at 0 hpi, 24 hpi, 48 hpi and 72 hpi in RenSeq dataset, whereas only 377.75, 355.5, 291 and 237.25 NLRs (average of 4 replicates) were 100% covered with RNAseq reads (Table 7). Similar to the expression data, cDNA enrichment sequencing provides a more robust analysis of the transcription across larger parts of the genes, including at early infection time points.

**Table 6:** Comparison between RNAseq and enrichment techniques for coverage of pathogen effector genes (RxLRs and non-RxLRs). Average was taken for 4 replicates.

| Hours post infection (hpi) | Average number of effectors with coverage above 0 |  | Average number of effectors with 100% coverage |  |
| --- | --- | --- | --- | --- |
|  | PenSeq | RNAseq | PenSeq | RNAseq |
| 0 | 288.75 | 146.25 | 99.00 | 8.75 |
| 24 | 331.75 | 181.25 | 132.25 | 15.25 |
| 48 | 331.00 | 192.00 | 100.25 | 19.25 |
| 72 | 311.50 | 149.50 | 58.00 | 12.25 |

**Table 7:**
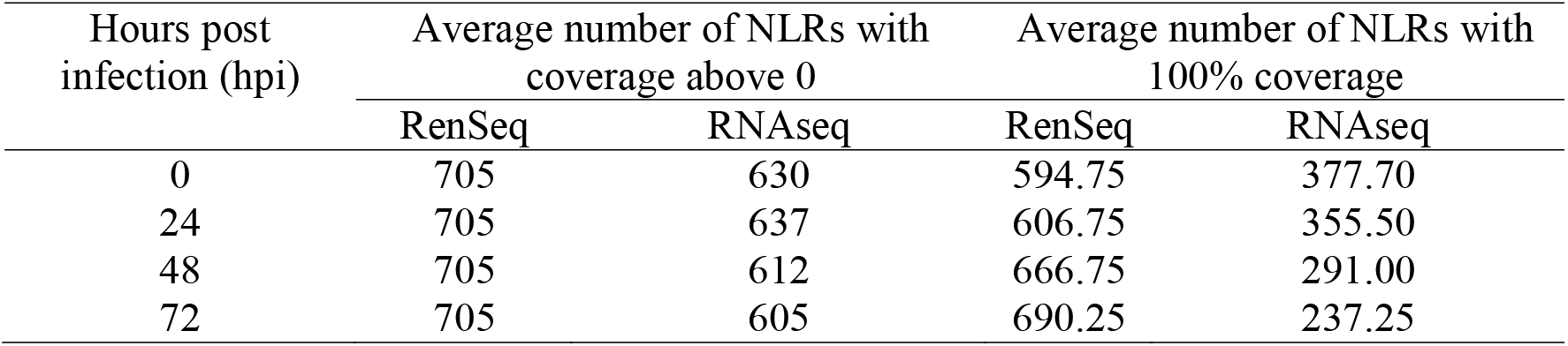
Comparison between RNAseq and enrichment techniques for coverage of NLRs. Coverage values for NLRs were calculated using all the 755 reported NLRs identified in Jupe et al. 2013 (according to DM v4.03). Average was taken for 4 replicates.

## 4. Discussion

Transcriptomics is a powerful method to enhance our understanding of molecular events occurring during infection. Although parallel expression studies of the host and pathogen during disease progression have provided insights into plant-pathogen interactions, the abundance of host biomass, and thus expressed genes during the early phases of infection, often limit detection and expression profiling of pathogen genes [42,43]. To overcome this limitation, we introduce cDNA-based target enrichment sequencing, RenSeq and PenSeq, for parallel NLRome and effectorome transcriptome studies during potato late blight infection. The use of cDNA RenSeq and PenSeq has been described previously to aid the annotation of expressed host and pathogen genes [14,24]. Here, we further assess if cDNA enrichment technologies enable quantitative transcriptomic analyses at early and late infection time-points.

Targeted enrichment sequencing in the form of RenSeq and PenSeq is facilitated through an excess of RNA-derived probes that drive the hybridisation to target sequences, which are represented through cDNA in this study. It is this molar excess of baits that is key to retaining the quantitative nature in the gene expression profiles post enrichment sequencing. Our comparative analysis between PenSeq and RNAseq reflects the hemibiotrophic nature of the infection. Previously utilised marker genes for the late blight infection progression such as *HMP1*, *NPP1* and known *Avr* genes [39,40,44] with a biotrophic expression profile, revealed concordance in their transcriptional behaviour for both RNAseq and PenSeq analyses (Figure 3, Supplementary Figure 3, Table S2).

In terms of biology, PenSeq and RNAseq revealed that the progression of infection in the tissue culture plant DM is very rapid, which is in line with the highly susceptible phenotype of DM and the lack of functional NLRs to control late blight [45]. This susceptibility is further promoted by the use of tissue culture derived plants. Consequently, in our replicated experiments, the visual transition from biotrophy to necrotrophy occurred at approximately 48 hours. No significant disease symptoms were observable at 24 hpi whereas later time points at 48 hpi and 72 hpi revealed progressive tissue collapse until complete plant death at 96 hpi (Figure 1). In line with this, cDNA PenSeq and RNAseq transcriptomic analyses showed that during the biotrophic phase the expression of the Haustorial Membrane Protein (*HMP1*; PITG_00375) was highest at 24 hpi (Supplementary Figure 3; Table S2). Similarly, *NPP1* (PITG_00957), a necrotrophic marker which shows the transition between the biotrophic and necrotrophic phases of infection [40], was highest at 48 hpi and after that it reduced significantly. This expression profile is in line with the observation made by [40], where in infections of tomato the expression of *NPP1* decreased after 48 hpi.

The high-level of expression correlation for transcripts that are detectable by RenSeq, PenSeq and RNAseq provides strong evidence that quantitative information is retained through the enrichment sequencing process. The correlation between RNAseq and enrichment datasets ranged between r = 0.78 —0.87, for PenSeq and r = 0.62—0.86 for RenSeq, both within and between replicates (Supplementary Figure 2, Table S4). The hierarchical clustering analysis further confirmed the similarity of expression patterns between PenSeq, RenSeq, and RNAseq data, while highlighting genes with no or low expression in the latter (Figure 3). Importantly, any gene identified as being expressed by the RNAseq analysis was also identified in the RenSeq or PenSeq analyses. However, RenSeq and PenSeq identified additional host and pathogen genes, respectively, that were below the threshold of expression detection in the RNAseq analysis. Notably, PenSeq was able to identify an average of 316 expressed genes throughout the time course, which is a two-fold increase compared to RNAseq, which identifies an average of 167 expressed genes only (Figure 2A, Table 2).

Targeted enrichment sequencing ensures minimal off-target sequencing and provides sufficient read depth and sequence coverage for identification of low expressed target genes [46]. In the present study, we observed that 89.98% of RNAseq reads and 76.99% of RenSeq reads map to the potato reference transcriptome. For PenSeq, averaged across all time points, an almost twelve-fold increase in the percentage of reads mapped to the *P. infestans* reference genome was observed in comparison with RNAseq. It is noteworthy that whilst in the RNAseq dataset, the reads mapped to the reference transcriptome remains almost constant throughout the infection time points at 0.2%, an increase in mapped reads from 1.04% at 0 hpi to 4.04% at 72 hpi was observed in PenSeq dataset (Table 1). This demonstrates the ability of PenSeq to capture transcripts from the pathogen during infection which enables the identification and quantification of expressed genes, many of which elude detection by RNAseq. In addition, the percentage coverage of targeted genes was more comprehensive for RenSeq and PenSeq compared to RNAseq, achieving complete coverage for about 80% of NLRs and 40% of effectors expressed at 24 hpi (Supplementary Figure 4).

Although limited to targeted genes of interest, our study highlights the advantages of cDNA-based enrichment sequencing techniques, PenSeq and RenSeq compared to RNAseq, for not only ascertaining gene expression, but also their transcriptional quantification. Specifically, these techniques offer insight into the detection and transcriptional profile of low expression genes with increased transcript coverage.

## CRediT authorship contribution statement

**Amanpreet Kaur:** Investigation, Data analysis, Visualization, writing-original draft. **Vikrant Singh:** Investigation, Data analysis, writing-original draft. **Stephen Byrne:** Data analysis. **Miles Armstrong:** Investigation, Methodology. **Brian Harrower:** Investigation, methodology. **Thomas Adams:** Review and editing manuscript. **Ewen Mullins:** Supervision, Project administration, Funding acquisition, Writing-reviewing & editing. **Ingo Hein:** Conceptualization, Supervision, Resources, Project administration, Funding acquisition, Writing-reviewing & editing.

## Acknowledgements

The authors acknowledge the Research/Scientific Computing teams at The James Hutton Institute and NIAB for providing computational resources and technical support for the “UK’s Crop Diversity Bioinformatics HPC” (BBSRC grant BB/S019669/1), use of which has contributed to the results reported within this paper.

## Sequence data availability

The RNAseq, RenSeq and PenSeq sequencing data derived from *Solanum tuberosum* group Phureja DM 1-3 516 R44 microshoots infected with *P. infestans* isolate W9928C are available at the European Nucleotide Archive (ENA) at EMBL-EBI as project PRJEB64683.

## Supplementary material

**Supplementary Figure 1:** Pipeline for cDNA pathogen-enrichment sequencing, resistance gene enrichment sequencing and RNAseq. RNA was extracted from leaves excised from *in vitro* plants after 0, 24, 48, 72, and 96 hours post infection (hpi) and processed for RNAseq, PenSeq and RenSeq in a parallel setup. The reads were mapped to reference genomes of *Solanum tuberosum* group Phureja DM 1-3 516 R44-v6.1 (DM v6.1) and *Phytophthora infestans* (strain T30-4) and the expression levels of effectors and NLRs were calculated and compared.

**Supplementary Figure 2** Correlation between replicates (average of 4 replicates) within and between **A)** PenSeq and RNAseq; **B)** RenSeq and RNAseq. Correlations between expression levels (TPM) of **C)** effectors in PenSeq and RNAseq datasets and **D)** NLRs in RenSeq and RNAseq datasets. The correlations were calculated as Pearson’s correlation coefficients. TPM values were normalized to log10(TPM+1) before plotting.

**Supplementary Figure 3:** Expression profiles of infection initiation through appressorium formation (cellulose synthase *CesA4*), biotrophy (Haustorium Membrane Protein; *HMP1*) and necrotrophy (Necrosis inducing Phytophthora Protein; *NPP1* like) marker genes as detected by PenSeq and RNAseq during different infection time points. TPM values were normalized to log10(TPM+1) before plotting.

**Supplementary Figure 4:** A comparison between RNAseq and enrichment sequencing techniques (PenSeq and RenSeq) for coverage and Read depth of effectors and NLRs. RNAseq, PenSeq and RenSeq reads were mapped to *Solanum tuberosum* group Phureja DM 1-3 516 R44-v4.03 (DM v4.03) and *P. infestans* T30-4 genome assembly. The coverages and read depth were calculated with bedtools [47].

**Table S1:** Number of reads from enrichment sequencing (PenSeq and RenSeq) and RNAseq passing the quality control parameters (minimum length 100 bp and phred score 20). The adaptors were trimmed using fastp in default settings.

**Table S2:** Expression values (Transcripts Per Million; TPM) of effectors (RxLRs and non-RxLRs) across different infection points as determined using PenSeq and RNAseq data

**Table S3:** Expression values (Transcripts Per Million; TPM) of NLRs across different infection points as determined using RenSeq and RNAseq data. Expression was calculated only for the NLRs for which reference gene IDs are available in the latest potato transcriptome version (v6.1)

**Table S4:** Correlation between RNAseq and PenSeq/RenSeq enrichment sequencing datasets and time points using Pearson’s correlation. Before analysing the Transcripts Per Million (TPM) values were transformed using log10 (TPM+1)

**Table S5:** Differential expression of effectors. A gene was considered significantly differentially expressed if it had an adjusted p-value of < 0.01 and log2 fold change ≥ 1

**Table S6:** Differential expression of NLRs. A gene was considered significantly differentially expressed if it had an adjusted p-value of < 0.01 and log2 fold change ≥ 1

