## Supplementary figures for "Transcriptional profiling during infection of potato NLRs and *Phytophthora infestans* effectors using cDNA enrichment sequencing"

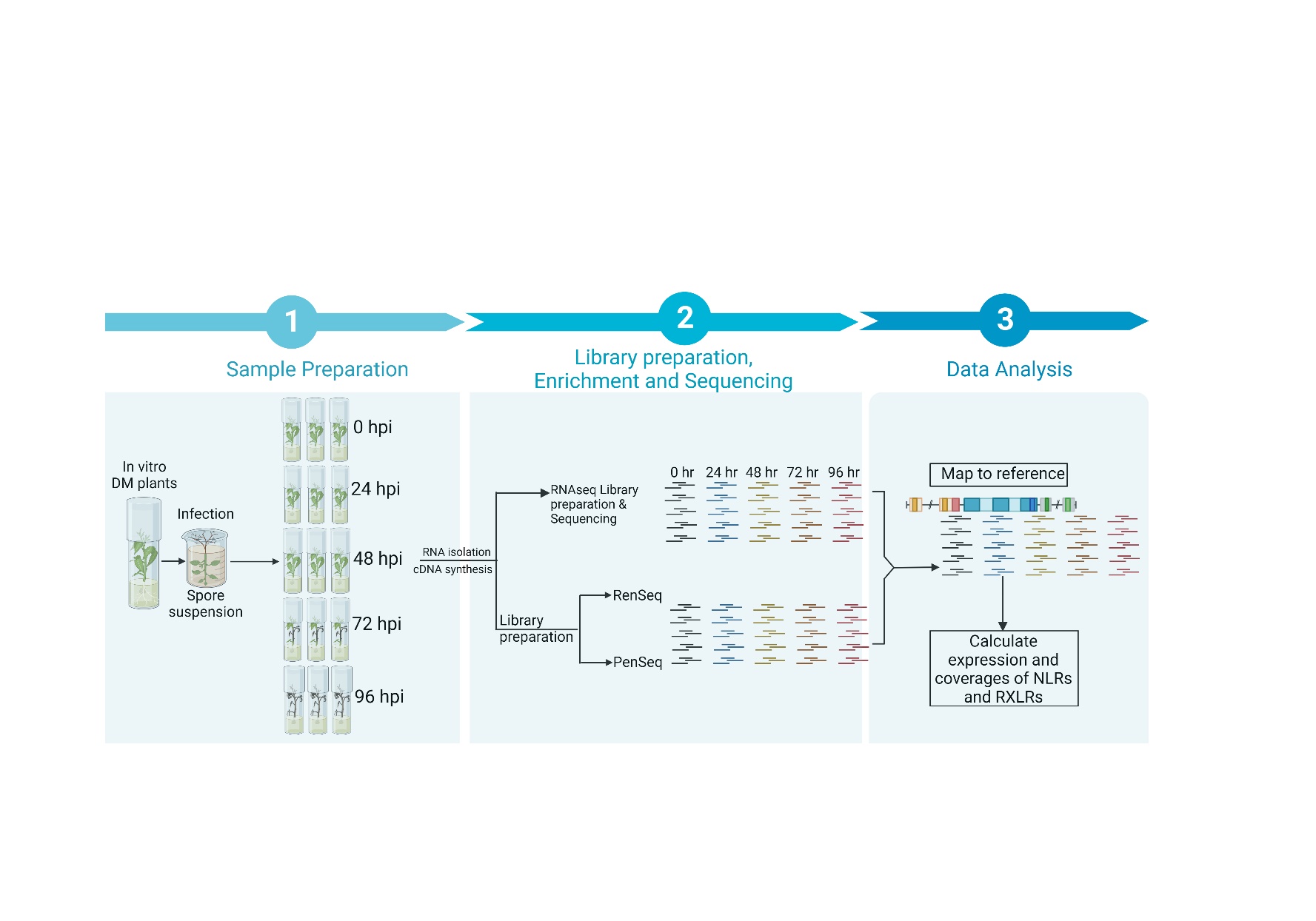

**Supplementary Figure 1:** Pipeline for cDNA pathogen-enrichment sequencing, resistance gene enrichment sequencing and RNAseq. RNA was extracted from leaves excised from *in vitro* plants after 0, 24, 48, 72, and 96 hours post infection (hpi) and processed for RNAseq, PenSeq and RenSeq in a parallel setup. The reads were mapped to reference genomes of *Solanum tuberosum* group Phureja DM 1-3 516 R44-v6.1 (DM v6.1) and *Phytophthora infestans* (strain T30-4) and the expression levels of effectors and NLRs were calculated and compared.

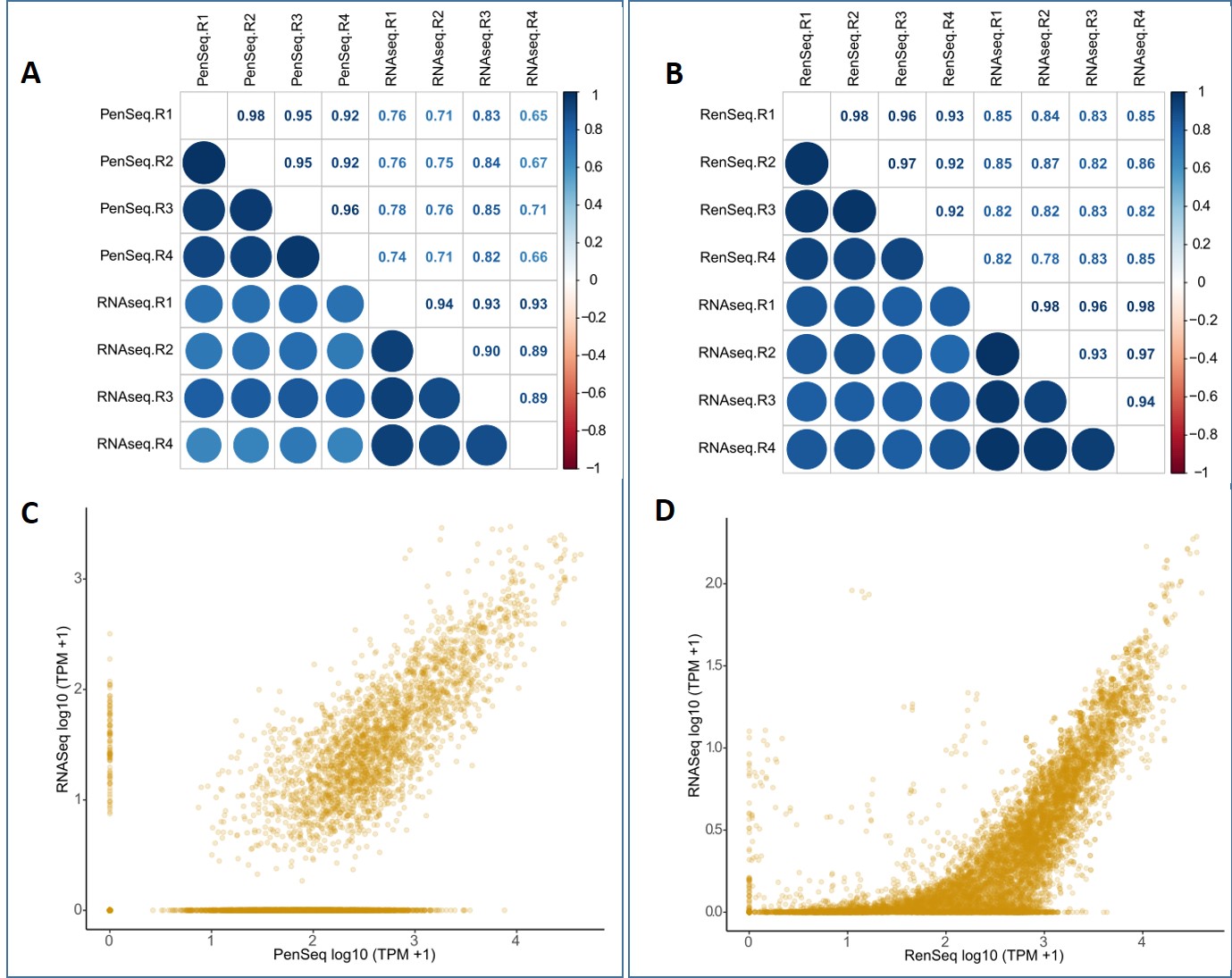

**Supplementary figure 2** Correlation between replicates (average of 4 replicates) within and between **A)** PenSeq and RNAseq; **B)** RenSeq and RNAseq. Correlations between expression levels (TPM) of **C)** effectors in PenSeq and RNAseq datasets and **D)** NLRs in RenSeq and RNAseq datasets. The correlations were calculated as Pearson’s correlation coefficients. TPM values were normalized to log10(TPM+1) before plotting.

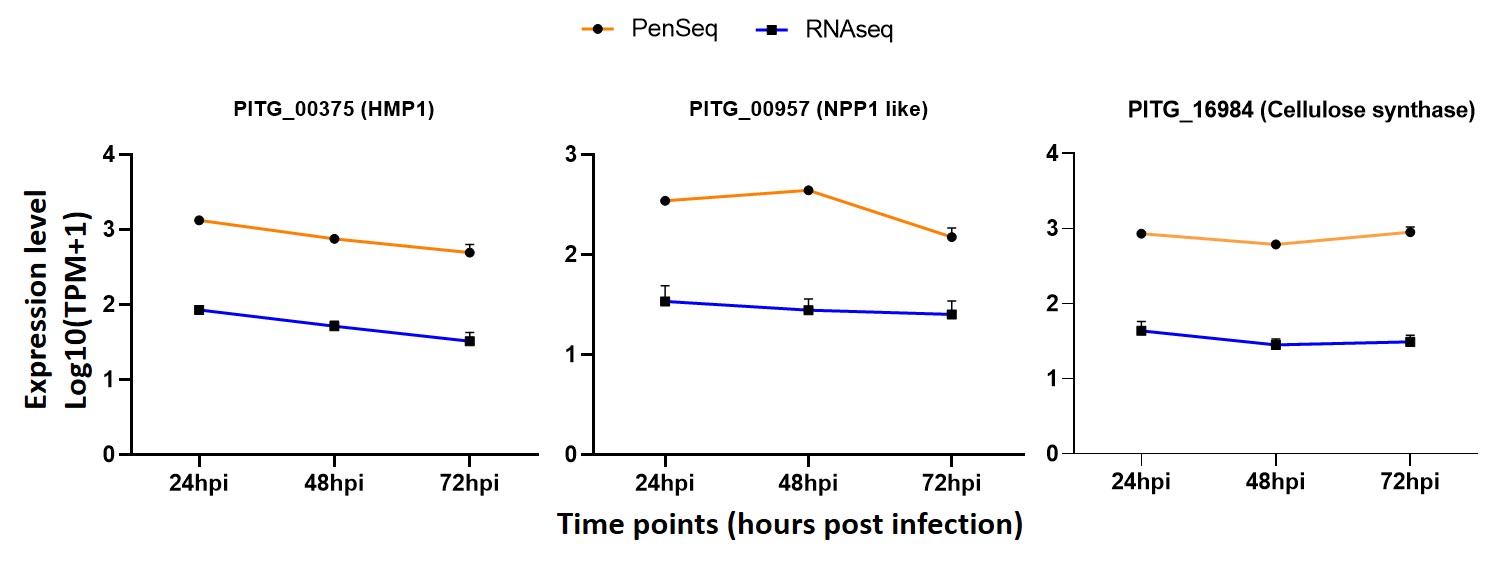

**Supplementary figure 3:** Expression profiles of infection initiation through appressorium formation (cellulose synthase *CesA4*), biotrophy (Haustorium Membrane Protein; *HMP1*) and necrotrophy (Necrosis inducing Phytophthora Protein; *NPP1* like) marker genes as detected by PenSeq and RNAseq during different infection time points. TPM values were normalized to log10(TPM+1) before plotting.

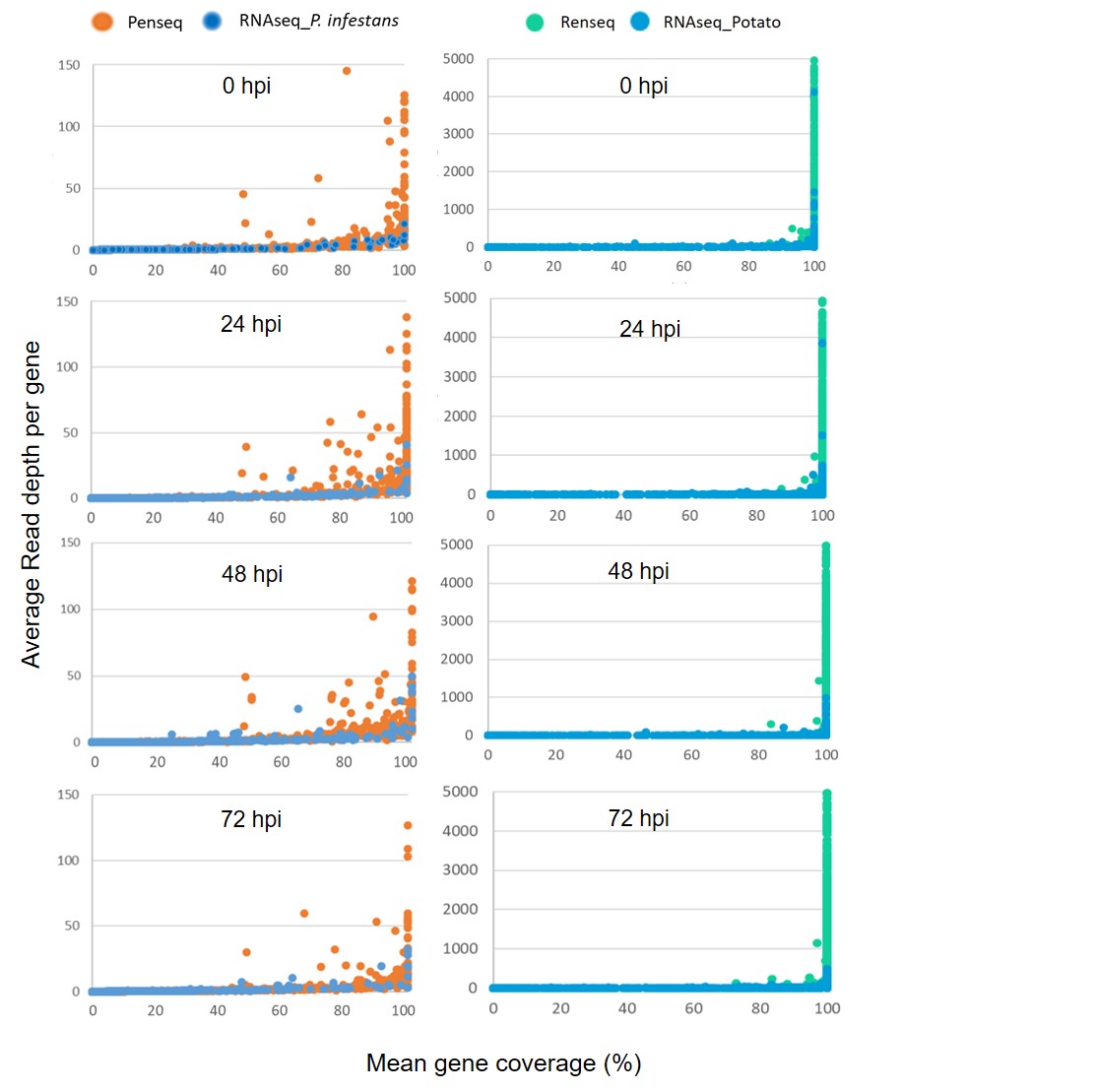

**Supplementary Figure 4:** A comparison between RNAseq and enrichment sequencing techniques (PenSeq and RenSeq) for coverage and Read depth of effectors and NLRs. RNAseq, PenSeq and RenSeq reads were mapped to *Solanum tuberosum* group Phureja DM 1-3 516 R44-v4.03 (DM v4.03) and *P. infestans* T30-4 genome assembly. The coverages and read depth were calculated with bedtools [47].
